# Microvessel Chaste: An Open Library for Spatial Modelling of Vascularized Tissues

**DOI:** 10.1101/105692

**Authors:** J.A. Grogan, A.J. Connor, B. Markelc, R.J. Muschel, P.K. Maini, H.M. Byrne, J.M. Pitt-Francis

## Abstract

Spatial models of vascularized tissues are widely used in computational physiology, to study for example, tumour growth, angiogenesis, osteogenesis, coronary perfusion and oxygen delivery. Composition of such models is time-consuming, with many researchers writing custom software for this purpose. Recent advances in imaging have produced detailed three-dimensional (3D) datasets of vascularized tissues at the scale of individual cells. To fully exploit such data there is an increasing need for software that allows user-friendly composition of efficient, 3D models of vascularized tissue growth, and comparison of predictions with *in vivo* or *in vitro* experiments and other models. Microvessel Chaste is a new open-source library for building spatial models of vascularized tissue growth. It can be used to simulate vessel growth and adaptation in response to mechanical and chemical stimuli, intra- and extra-vascular transport of nutrient, growth factor and drugs, and cell proliferation in complex 3D geometries. The library provides a comprehensive Python interface to solvers implemented in C++, allowing user-friendly model composition, and integration with experimental data. Such integration is facilitated by interoperability with a growing collection of scientific Python software for image processing, statistical analysis, model annotation and visualization. The library is available under an open-source Berkeley Software Distribution (BSD) licence at https://jmsgrogan.github.io/MicrovesselChaste. This article links to two reproducible example problems, showing how the library can be used to model tumour growth and angiogenesis with realistic vessel networks.

## Introduction

Spatial models of vascularized tissue are used to study the growth and response to treatment of tumours [1], angiogenesis [2], osteogenesis [3], coronary perfusion [4] and tissue oxygenation [5]. Such models typically comprise a combination of: i) agent-based or continuum representations of migrating and proliferating cells, ii) line-based or spatially resolved representations of microvessels, iii) the solution of blood, nutrient, growth factor and drug transport problems in vessel networks whose geometry and connectivity may evolve, iv) the solution of growth factor and drug transport problems in the evolving extravascular space and v) vessel formation and endothelial tip cell migration in response to mechanical and chemical cues.

Composition of spatial models of vascularized tissue growth is a time consuming process, with most researchers writing custom software. There are many examples of such models, which have typically been implemented using MATLAB, C, C++ and Fortran, including Anderson and Chaplain [6], Alarcón *et al.* [7], Friboes *et al.* [8], Shirinifard *et al.* [9], Owen *et al.* [1], Perfahl *et al.* [10], Welter and Rieger [11], Secomb *et al.* [2], and Boas and Merks [12]. The need to account for many biological phenomena has made the development of more general software frameworks challenging [13,14], with functionality for integration with experimental imaging data and model benchmarking and cross-comparison also important [14,15]. Several groups, including Popel and co-workers [16], Secomb and co-workers [2] and Beard, Bassingthwaighte and co-workers [17], have produced software that focuses on general modelling and integration with imaging data. Notable efforts in this area are being undertaken as part of the European Union's Virtual Physiological Human Project (http://www.vph-institute.org/) and the National Institute of General Medical Sciences’ Virtual Physiological Rat (http://www.virtualrat.org/) program.

Microvessel Chaste is a new open-source Python/C++ library for composing spatial models of vascularized tissues. It is a ‘plug-in’ for the Chaste C++ library for problems in computational physiology and biology [18], integrating with existing cell-based and sub-cellular models and partial differential equation (PDE) and ordinary differential equation (ODE) solvers. Development has been motivated by the above mentioned vascularized tissue studies and software. However, there is an additional focus on providing a user-friendly, general framework for model composition, in a manner similar to that in which Chaste [18], CompuCell3D [19], EPISIM [20] and PhysiCell [21] can be used to compose tissue models with agent-based representations of cells. The primary strength of Microvessel Chaste is that it provides a comprehensive Python interface. This facilitates integration with a growing collection of scientific Python software for image processing (scikit-image (http://scikit-image.org/), ITK (https://itk.org/), VMTK (https://vmtk.org/), statistical analysis (SciPy (https://www.scipy.org/), pandas (http://pandas.pydata.org/)), model annotation (libSBML (http://sbml.org/Software/libSBML)) and visualization (matplotlib (http://matplotlib.org/), VTK [22]). It allows easy extension by the user, who can compose their own readers, writers and solvers without re-compiling the code. The library has facilitated the integration of modelling with high resolution three-dimensional (3D) imaging data, as shown in Grogan *et al.* [23] and below, and will be useful in future model cross-comparison studies.

The next section overviews the library design and implementation. This is followed by example simulations of tumour growth, with a realistic vessel network, and a corneal micropocket angiogenesis assay.

## Materials and Methods

This section summarizes available algorithms by demonstrating how the library can be used to construct a typical multi-scale tissue growth simulation. The library can be installed on Linux using the Conda package manager (http://conda.pydata.org/), and used in a Jupyter notebook (http://jupyter.org/). Alternatively, it can be built from source using CMake. Docker images with a running Jupyter notebook server are available for Windows and MacOS.

### Capabilities and implementation

Fig. 1 shows the stages involved in setting up a detailed multi-scale vascularized tissue growth simulation using Microvessel Chaste. Simulations can be constructed in a flexible manner thanks to the use of object-oriented programming. In this example, a simulation domain, vessel network and cell population are first constructed. Cell and vessel locations and attributes can be described using NumPy arrays (http://www.numpy.org/) and are automatically copied to and from uBLAS vectors when interfacing with C++ routines. The user can easily implement their own vessel network reading and writing modules in Python, which is useful as there is no standard format for describing vessel network coordinates and connectivity. PDEs for transport of diffusible chemicals through the tissue can be configured, and rules for vessel growth or shrinkage due to blood flow defined, along with rules for vessel sprouting and endothelial tip cell migration. Custom PDE solvers can be used, with solutions returned to other solvers at requested sample points through C++ or Python interfaces.

**Figure 1.**
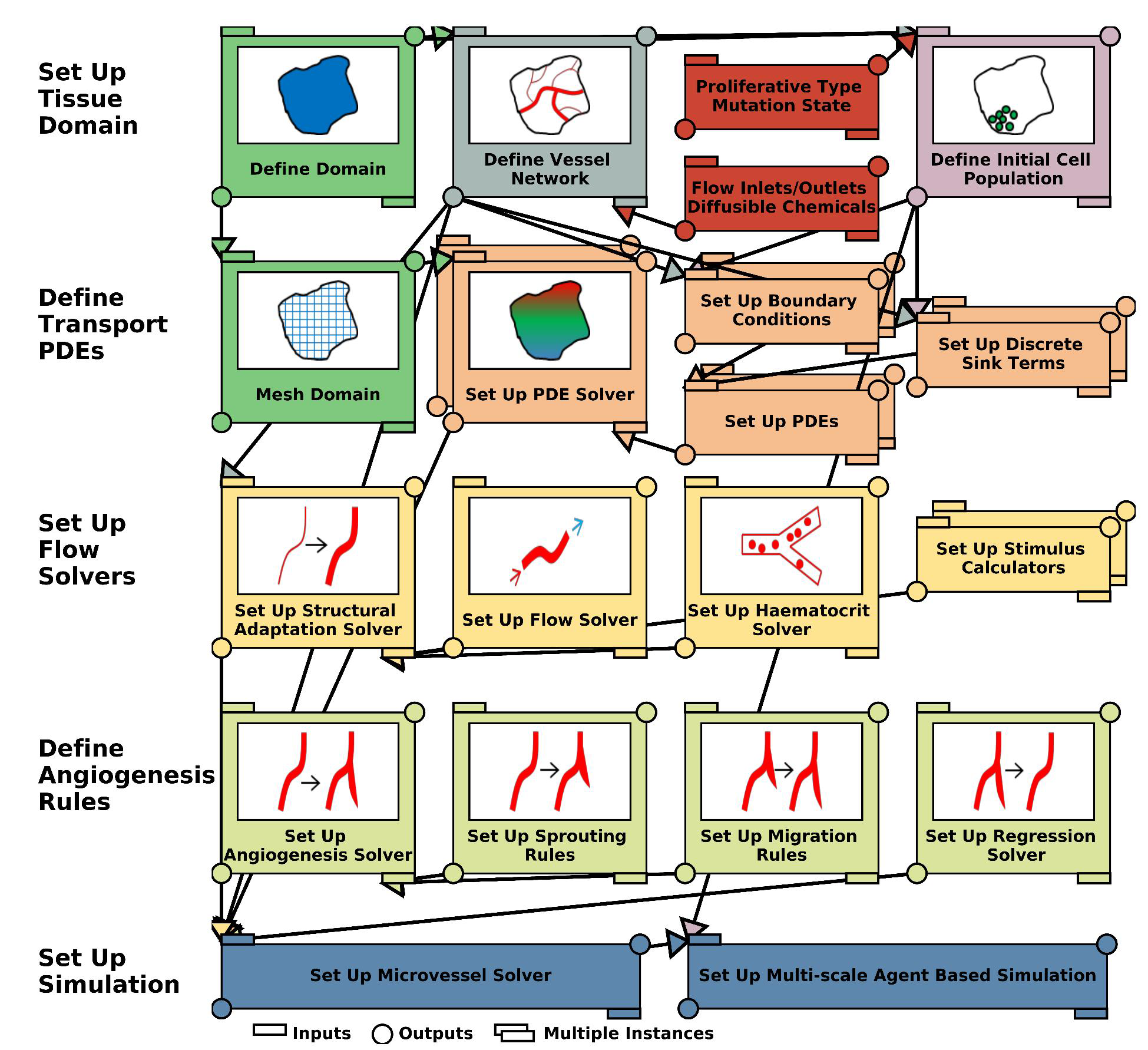
A schematic showing the composition of a detailed multi-scale vascularized tissue growth model using Microvessel Chaste. Multiple instances of certain components can be used, for example to define a series of sequentially coupled PDEs.

The domain, which is a geometrical feature (specifically a piecewise linear complex [24]) in two dimensions (2D) or 3D, can be used to construct vessel networks and cell populations through multiple space-filling or Boolean operations, and to generate a computational grid (mesh) for use in the solution of PDEs. Geometries and meshes can be read and written in VTK or STereoLithography (STL) formats. Sub-models can be collected in hierarchical structures, for example a `StructuralAdaptationSolver` can manage a `FlowSolver` and is itself managed by a `MicrovesselSolver`. Alternatively, sub-models can be executed in isolation, for example to solve a nutrient PDE with cell location dependent sink terms.

The model can be solved using a suitable time-stepping scheme, available in Chaste. During a time step, nutrient and angiogenic stimulus concentration fields can be updated. Cells can migrate and progress through their cycle, and vessels can form sprouts, regress or alter their radii. Cell and vessel data and PDE solutions can be visualized using built-in VTK-based rendering, output in VTK format for visualization in Paraview, or returned as NumPy arrays for further processing.

### Code layout and design

The components of the library are as follows:

- `geometry:` generation of 2D and 3D piecewise linear complex geometries for direct use with Tetgen [24] meshing software;
- `mesh:` automated finite element meshing of 2D and 3D geometries and interpolation of vessel and cell locations onto unstructured and structured grids;
- `ode:` ODE models for progress through the cell cycle;
- `pde:` PETSc [25] based finite difference and finite element solvers for steady-state reaction-diffusion equations with discrete sinks and sources at vessels and cells;
- `population:` analysing, reading, writing and generating vessel networks and cell populations;
- `simulation:` flow, structural adaptation and angiogenesis solvers. Simulation modifiers for integration with discrete cell simulations in Chaste;
- `utility:` dimensional analysis, a collection of literature parameters of interest for vascularized tissue simulations and 3D visualization tools.

Object-oriented programming is used throughout. Python bindings are generated automatically using Py++, exposing a significant amount of C++ functionality. Low-cost, compile-time, unit checking is used to ensure dimensional consistency through the Boost Units framework [26]. The library includes detailed API documentation (full Doxygen coverage) and includes over 80 C++ and Python unit test suites for individual components. Several reproducible C++ and Python (Jupyter notebook) tutorials are available, including for the two example problems in the next section.

## Results

In this section tumour growth and angiogenesis problems are demonstrated; they are available for reproduction at https://jmsgrogan.github.io/MicrovesselChaste. A collection of additional, simpler examples is also available at this location.

### A 3D tumour growth simulation

The first example, shown in Fig. 2, is a 3D simulation of tumour growth in a vessel network geometry obtained using multi-photon imaging after implantation of MC38 tumour cells in a mouse [23]. The tumour growth model is similar to many in the literature [6, 8, 10, 27]. However, the use of a large, realistic and evolving tumour vessel network distinguishes this example from previous studies. The simulation is facilitated by recent advances in intravital imaging, which allow *in vivo* observation of tumour growth at the scale of individual cells, and the new pre-processing and modelling functionality in Microvessel Chaste.

**Figure 2.**
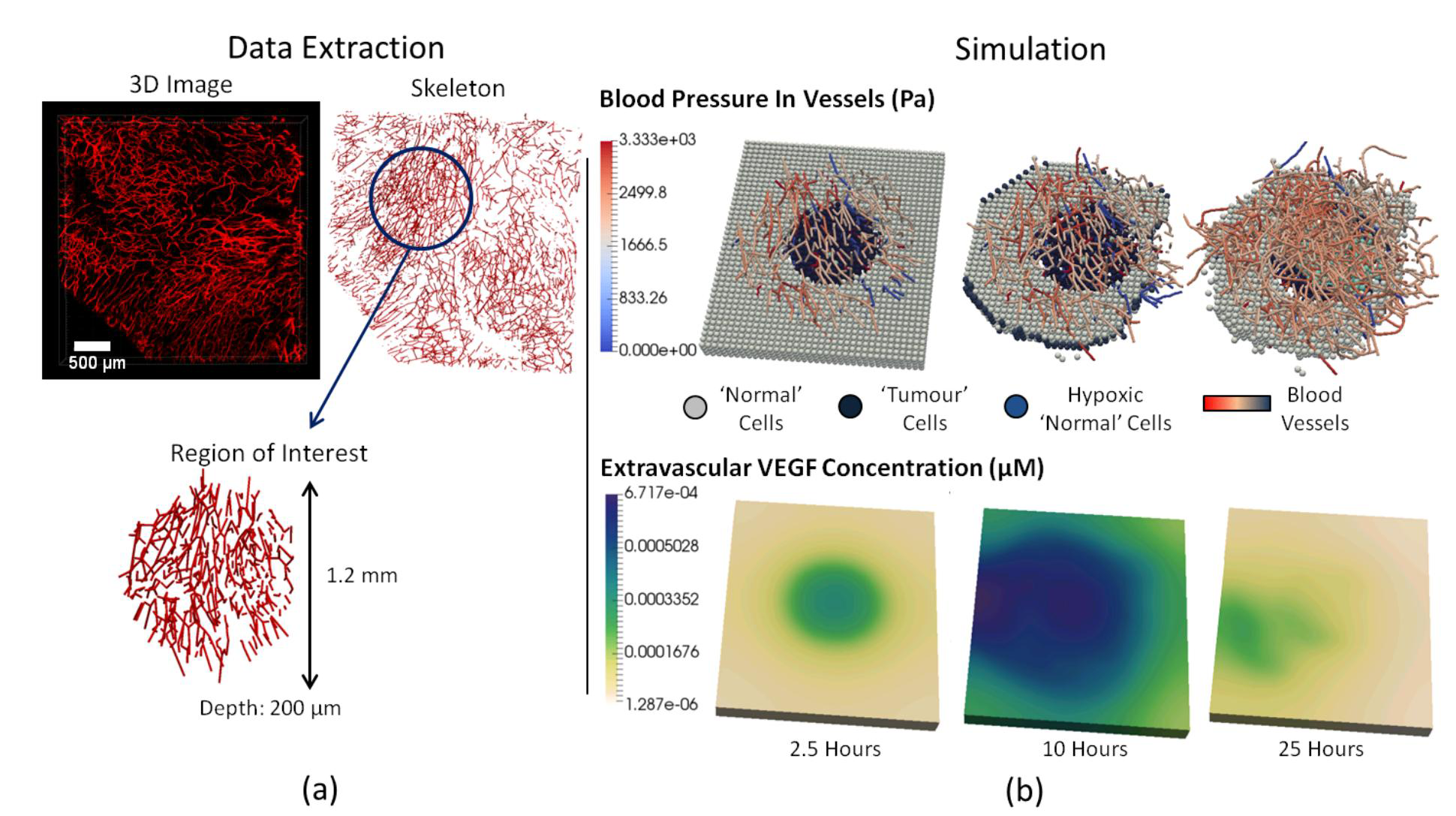
(a) A 3D intravital image of a tumour microvessel network (red) is obtained and a skeleton extracted as described in Grogan *et al.* [23]. A cylindrical region of interest with diameter 1.2 mm is extracted for the example simulation. (b) A tumour growth simulation using the extracted microvessel network and Microvessel Chaste. The predicted evolution of the tumour over 25 hours is shown, including blood pressure in growing vessels, VEGF concentrations in the extravascular space and discrete cells.

The problem is initialized with a 3D region of the tumour vessel network. A regular lattice is generated in the bounding box of the network geometry. A cellular automaton based cell population fills all lattice sites, including those occupied by vessels. ‘Tumour’ cell types are assigned to a central cylindrical region and ‘Normal’ types to the remainder. Cells far from oxygen rich vessels experience low oxygen levels and, as a result, become hypoxic and release Vascular Endothelial Growth Factor (VEGF). VEGF stimulates the sprouting and chemotactic migration of new vessels from the existing vasculature. At later times, cells far from the vessels become apoptotic. The surviving tumour cells gradually invade the domain at the expense of the normal cells. This process is similar to those observed in the simulations of Perfahl *et al.* [10], Anderson and Chaplain [6] and others. There are many potential extensions to models of this type, including simulated administration of chemo-therapeutic and anti-angiogenic drugs [7] and radiotherapy [23]. These cases can be simulated using the Microvessel Chaste library.

### A 3D angiogenesis simulation in a curved domain

Our second example is a 3D, off-lattice simulation of angiogenesis in a curved geometry (see Fig. 3). This example demonstrates geometry manipulation and the solution of PDEs on 3D domains. The application is appropriate for the corneal micropocket assay that is widely used to study angiogenesis [27]. Typical experimental results are shown in Fig. 3(a).

**Figure 3.**
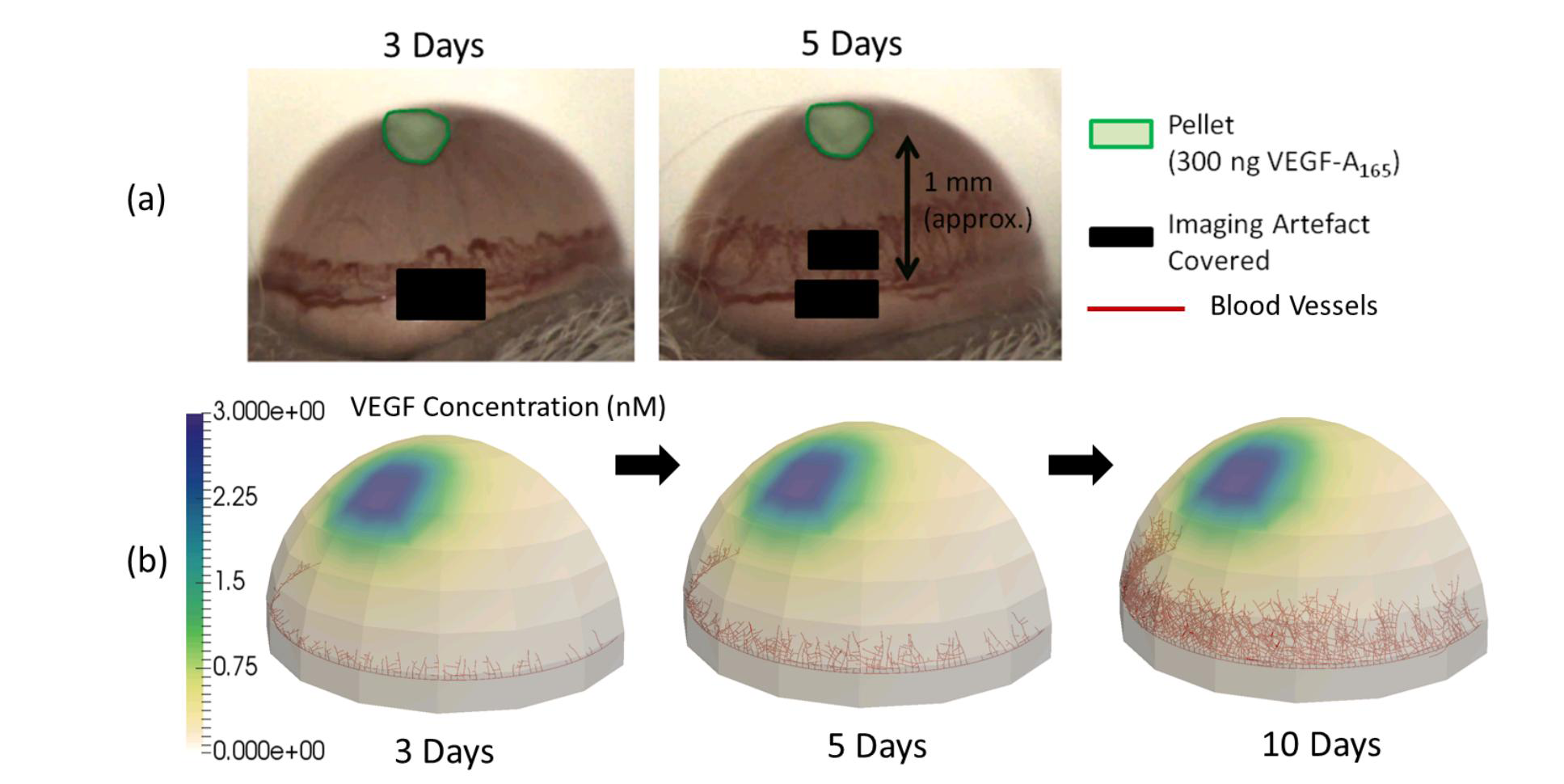
(a) Images from a cornea micropocket experiment showing microvessels (dark red) at 3-5 days post pellet implantation [27]. (b) Application of the Microvessel Chaste library in modelling a similar experiment.

In this experimental assay a pellet containing an angiogenic growth factor (for example, VEGF) is implanted in the cornea. VEGF diffuses from the pellet into the corneal tissue and stimulates endothelial cells lining existing vessels at the base to form sprouts. The sprouts then migrate toward the pellet along spatial gradients in VEGF. This example follows a common modelling paradigm where agent-based representations of cells are not included, but individual vessels are [2]. As shown in Fig. 3(b), the cornea is represented as a hemispherical domain of radius 1.4 mm and thickness 0.1 mm. The pellet is a cuboid with side length 0.3 mm and depth 0.1 mm, with a prescribed VEGF concentration of 3.0 nM on the boundaries. In practice, the VEGF in the pellet will deplete. Vessels sprout from the limbal vessel and then follow a persistent random walk with bias towards other vessel tips and up VEGF gradients. The VEGF distribution is obtained by solving a reaction-diffusion PDE on the cornea at the start of the simulation, and is fixed for the remainder. Vessels migrate toward the pellet, remaining within the volume of the 3D cornea geometry. Possible extensions to this simple model include the addition of discrete stromal cells, distinction between perfused and unperfused vessels, subcellular signalling, VEGF depletion and consumption by cells, and the use of multiple vessel growth factors, as per Connor *et al.* [27].

## Conclusions

A new library for composing spatial models of vascularized tissues has been demonstrated, and two reproducible sample problems in the areas of tumour growth and angiogenesis presented. Additional functionality for semi-automated 2D and 3D image segmentation and meshing is under development, to aid further integration with experimental studies such as those shown in Fig. 2(a). Porting to Windows and MacOS is also of interest. At present most algorithms operate in serial only, however all PDE and flow solvers are based on PETSc [25] structures and vessel network components may be communicated across processors using existing serialization functionality in Chaste [28]. The library is available under a permissive BSD license, with source files and documentation available via the project Github page https://jmsgrogan.github.io/MicrovesselChaste/. Contributions are welcome via Github pull requests and issues can be reported via the Github issue tracker. The latest release, version 3.4.2, is archived at doi.org/10.5281/zenodo.213148.

## Author Contributions

JAG, AJC, PKM, HMB and JMPF designed the models and software. JAG, AJC and JMPF developed the software. BM and RJM designed the experimental imaging. BM performed the experimental imaging. JAG, AJC, BM, PKM, HMB and JMPF drafted and edited the article. All authors read and approved the final article.

## Acknowledgments

The research leading to these results has received funding from the People Programme (Marie Curie Actions) of the European Unions Seventh Framework Programme (FP7/2007-2013) under REA grant agreement No 625631 (BM) and the European Union’s Seventh Framework Programme for research, technological development and demonstration under grant agreement No 600841 (JAG, AJC, HMN, PKM, JMPF). BM and RJM acknowledge that this work was also supported by Cancer Research UK (CRUK) grant number C5255/A18085, through the CRUK Oxford Centre, by CRUK grant number C5255/A15935 and by CRUK/EPSRC Oxford Cancer Imaging Centre (grant number C5255/A16466). The authors acknowledge helpful inputs from the Chaste development team, in particular Jonathan Cooper, Alex Fletcher, James Osborne, Gary Mirams and Martin Robinson.

